# The Hedgehog receptors PTCH1 and PTCH2 exist as active homomeric and heteromeric complexes

**DOI:** 10.1101/2023.08.08.549832

**Authors:** Alex J. Timmis, Felix Cross, Danai S. Gkotsi, Hattie Ollerton, Colin A. Johnson, Natalia A. Riobo-Del Galdo

## Abstract

With the importance of Hedgehog signalling in embryonic development, tissue homeostasis and disease, understanding the molecular mechanisms of signal transduction is paramount for the design of specific, effective therapeutics. The Hedgehog receptor PTCH1 and the less studied PTCH2 isoform are evolutionarily related to bacterial RND permeases and sterol sensing proteins, which mobilise hydrophobic compounds powered by a cation gradient. Here we demonstrate that, in the active state, PTCH1 and PTCH2 form homomeric and heteromeric complexes that are inhibited by binding of Sonic Hedgehog. We show that PTCH2, unlike PTCH1, appears to have minimal cholesterol transport activity, but that conserved residues involved in cation transport are essential for its function. Heteromeric PTCH1-PTCH2 complexes depend on PTCH1’s cholesterol transport capacity, but the cation transport can be provided in trans by PTCH2, suggesting that some deleterious mutations in either isoform can be silenced by formation of heteromers, enhancing the robustness of this signal transduction system. These findings provide the molecular basis for the intriguing behaviour of PTCH2 as semi-redundant and partially overlapping in function with PTCH1 and explain the dominant negative effect of mutations that disrupt the PTCH2 cation transport triad in rare cases of cancer.

## INTRODUCTION

The Hedgehog (Hh) signalling pathway has a central role during embryonic development and its dysregulation in mature organisms is associated with disease states (1, 2). Hh signalling is activated by binding of one of three secreted Hh glycoproteins, Sonic Hh (Shh), Indian Hh (Ihh) or Desert Hh (Dhh) to their receptors Patched1 (PTCH1) and the less studied Patched2 (PTCH2) (3). Unlike most receptors, in the absence of ligand Patched proteins actively repress another membrane protein called Smoothened (Smo), a GPCR that mediates activation of the Gli family of transcription factors which drive canonical Hh signalling. Loss of function of PTCH1 leads to constitutive cell-autonomous activation of Smo/Gli signalling, underlying cancer development.

The Patched proteins topology consists of 12-transmembrane domains, intracellular N- and C-termini, a large intracellular middle loop and two extracellular domains which interact with the Hh ligands. Recent advances in structural biology have allowed the investigation of PTCH1 by cryo-electron microscopy (cryo-EM). Purified PTCH1 proteins with partly deleted middle loop and C-terminal domains were described in monomeric and dimeric form (4). The cryo-EM structures also showed the presence of several sterol molecules in an apparent hydrophobic tunnel, suggesting that PTCH1 is a sterol transporter, a biochemical activity that was later demonstrated (4–6). Co-purification of receptor ligand complex showed that the inhibited receptor exists as a 2 PTCH1:1 Shh complex where PTCH1 exists as an asymmetric dimer or “dimer of dimers” (7, 8). The complex is bridged by a molecule of Shh, which in its physiological mature form is a dually-lipidated glycoprotein, here abbreviated pN-ShhChol (reviewed in 9). The N-terminus of all three Hh proteins are palmitoylated by Hedgehog acyl transferase (HHAT) following an autocatalytic cleavage induced by formation of a cholesterol ester bond to the carboxyl group of the newly formed C-terminus. Fully modified Shh is highly asymmetric, with an extended N-terminal palmitate arm and a globular Ca^2+^-binding domain. In the inhibited state, pN-ShhChol binds to one PTCH1 monomer through its palmitate N-terminal modification and to the other monomer through an interphase of the globular Ca^2+^-binding domain by a pincer grasp mechanism (8). The palmitoylated N-terminus of Shh occupies part of the sterol hydrophobic tunnel, and a short palmitoylated N-terminal fragment of Shh (pShh22) is sufficient to inhibit PTCH1 activity and stimulate canonical Hh signalling (10). Similar to bacterial RND permeases, PTCH1 cholesterol transport requires powering by a cation gradient, which is likely transported in the opposite direction of the sterol. Several independent studies propose that the K^+^ gradient is essential for PTCH1 activity (11, 12), although others provided evidence for a role of the Na^+^ gradient (13). Regardless of the cation identity, a conserved GxxxDD motif in TM-4 is essential for PTCH1 activity, that is its capacity to inhibit Smo-dependent Gli activation.

PTCH1 is an essential gene necessary for preventing ectopic cell autonomous, ligand-independent activation of the canonical Hh pathway. Mice deficient in *Ptch1* die *in utero* with a phenotype of overgrowth, CNS and limb patterning defects and more subtle alterations in most tissues’ architecture consistent with hyperactivation of Hh signalling (14). In contrast, mice deficient in *Ptch2* have a much more subtle phenotype and live to adulthood (15). A few studies suggested a partially overlapping role of PTCH2 with PTCH1 in skin and limb development (16, 17). Furthermore, Ptch2 was shown to mediate a small response to Shh in *Ptch1^−/−^* mouse embryonic fibroblasts and embryonic bodies (18). A clearer picture of the role of PTCH2 as a positive modulator of PTCH1 activity was evident in conditions of haploinsufficiency of *Ptch1*, where deficiency of Ptch2 exacerbates tumorigenesis (15). Altogether, PTCH2 seems to have SMO repressive activity and to respond to Shh, but its function is eclipsed in the presence of PTCH1.

In this study, we characterised the oligomeric state of active (ligand-free) PTCH1 and PTCH2 in their natural environment in live cells by a combination of Förster resonance energy transfer (FRET)-reduction after photobleaching, co-immunoprecipitations, and Gli-reporter activity assays. Our findings reveal that, in agreement to the cryo-EM structures of PTCH1, both PTCH1 and PTCH2 can form homomeric complexes in live cells in the absence of a bridging ligand. Furthermore, we found that PTCH1 and PTCH2 can exist as active complexes with an intermediate potency to inhibit SMO between PTCH1 homomers (highest activity) and PTCH2 homomers (lowest activity). Remarkably, the activity of heteromeric complexes seems to depend on the cholesterol transport activity of the PTCH1 monomer but that cation antiporter can be provided in *trans* by PTCH2. These findings explain the partially overlapping function of PTCH1 and PTCH2 in developing tissues and postnatal homeostasis and provide a unifying model that explains some seemingly incongruent earlier reports.

## MATERIALS and METHODS

### Cell lines and culture procedures

HEK 293 human embryonic kidney epithelial cells and NIH 3T3 murine embryonic fibroblast cells were obtained from American Type Culture Collection (ATCC). *Ptch1^−/−^* mouse embryonic fibroblasts (MEFs) were a gift from Dr. Matthew Scott (Stanford University) and described in (19). Cells were maintained in Dulbecco’s Modified Eagle’s Medium (DMEM) (Gibco), supplemented with 10% Foetal Bovine Serum (FBS) (HEK 293) or Calf Serum (NIH 3T3 and *Ptch1^−/−^* MEFs) containing 1% Glutamax (Gibco). All cell lines were grown at 37°C in 5% CO_2_ and passaged prior to reaching confluence.

### Plasmids

HA-tagged mouse Ptch1 (Ptch1-HA) was a generous gift of Prof. Patrick Mehlen (Centre National de la Recherche Scientifique, CNRS, Lyon). Following generation of the Ptch1 K1413R-HA mutant by site directed mutagenesis (20), the HA-epitope was replaced by a 6xHis tag using the Q5 Site Directed Mutagenesis Kit (New England Biolabs) to generate Ptch1 KR-His. Additional rounds of site directed mutagenesis using the QuikChange XL II site directed mutagenesis kit (Agilent) were applied to create Ptch1 KR (ATT/LIN)-His, Ptch1 KR P490A-His, Ptch1 KR L504A-His and Ptch1 KR F1109A-His.

Enhanced green fluorescent protein (eGFP)-tagged human PTCH1 (PTCH1-eGFP) and myc-PTCH1 were a gift from Dr. Toshiyuki Miyashita (Kitasato University, Japan). The cDNA of eGFP was replaced by mCherry to generate PTCH1-mCherry using a two-step cloning strategy.

PTCH2-FLAG was created by a two-step cloning process, from the parental plasmid, pcDNA-PTCH2-pcDNA-HisB (gift from Prof. Peter Zaphiropoulos, Karolinska Institute). First, the coding region of PTCH2 was subcloned into the NotI and XbaI in pcDNA 3.1^+^. Next, a C-terminal FLAG tag ‘DYKDDDDK’ was introduced in frame, using InFusion cloning. PTCH2-FLAG was then fused N-terminally to eGFP or mCherry (PTCH2-eGFP and PTCH2-mCherry). PTCH2-HA was generated from PTCH2-FLAG by Q5 Site Directed Mutagenesis. PTCH2 LLW (L82F, L85F, W86A) and PTCH1 VLW (V111F, L114F, W115A) were generated in tandem by QuikChange XL II mutagenesis to generate PTCH2-LLW-FLAG, PTCH2-LLW-mCherry, PTCH1-VLW-eGFP and PTCH1-VLW-mCherry.

All mutants and fusion constructs were sequence verified.

### Gli-Luciferase Assay

*Ptch1^−/−^* MEFs were seeded in 24-well plates and transfected using TransIT-X2^®^ (Mirus) and Opti-MEM reduced serum media (Thermo Fisher Scientific). Briefly, each well was transfected with 50 μL mix containing 250 ng 8xGBS-Luc (a kind gift of Prof. Hiroshi Sasaki), 5 ng pRL-SV40 (Promega), and 375 ng total amount of testing plasmids individually or in combinations, according to the manufacturer’s instructions. In the cases where testing DNA concentrations did not total 375 ng, empty pcDNA 3.1^+^ plasmid DNA was added to make up the remaining quantity. Transfected cells were incubated for 24 h at 37°C, 5% CO_2_, before being carefully washed with 0.5 mL PBS per well. Media was then replaced with DMEM, 0.5% calf serum and 1% Glutamax. After 48 h, cells were washed with PBS and lysed with 100 μL of 1X passive lysis buffer per well. Lysis was performed at room temperature with shaking for 15 min. *Firefly*-Luciferase and *Renilla*-luciferase activities were determined using the Dual-Luciferase Reporter Assay System (Promega) in a Glomax 20/20 luminometer (Promega), as per manufacturer’s directions. The ratio *Firefly*-Luciferase/*Renilla*-luciferase measurements is presented as relative luciferase units, percent of control, of fold of control.

### Co-immunoprecipitation assays

HEK 293 cells were seeded at a density of 2 x 10^5^ cells/mL in growth medium without antibiotics and transfected 24 h later using Lipofectamine 2000 (Invitrogen). Briefly, 4 μg of each of two plasmids and 20 μL Lipofectamine 2000 were combined following the manufacturer’s directions for transfection of a 10-cm dish. After 24-36 h, the transfected cells were washed with ice-cold PBS and scraped in 700 μL Co-IP lysis buffer (50 mM Tris-HCL pH 7.5, 150 mM NaCl, 1% Nonidet P-40, 0.5% Sodium deoxycholate, 1 mM EDTA, 2.5 mM MgCl_2_ supplemented with 1x Proteoloc protease inhibitor, 0.4 mM PMSF, 1 mM DTT). Cell lysates were incubated 30 min at 4°C with rotation followed by centrifugation at 13,000 rpm at 4°C for 15 min. The supernatant was split into 200 μL for whole cell lysate and 500 μL was used for co-immunoprecipitation. The lysate was incubated with epitope tag-directed antibodies (Supplementary Table 1) for 1.5 h at 4°C with rotation, followed by addition of 30 μL Dynabeads (Invitrogen) and incubated for additional 2 h. The beads were washed 3 times with 1 mL Co-IP lysis buffer using a magnetic rack and the immunoprecipitates were eluted with 18 μL 2X Laemmli buffer (Pierce™) and heated at 45°C for 25 min. Whole cell lysates and co-IP samples were stored at −80°C for a maximum of 1 week before analysis.

**Supplementary Table 1.** Antibodies used for immunoblotting (WB), immunoprecipitation (IP) and immunofluorescence (IF) in this work. Abbreviations: CTS, Cell Signalling Technology; HRP, horseradish peroxidase.

| Antibody | Host species | Application | Dilution | Company | Catalog |
| --- | --- | --- | --- | --- | --- |
| **Arl13b** | Rabbit | IF | 1:500 | Proteintech | 17711-1-AP |
| **Myc (9B11)** | Mouse | IF | 1:500 | CST | 2276 |
| **Myc** | Mouse | IP | 1:2000 | Proteintech | 60003-2-Ig |
| **GFP (4B10)** | Mouse | IF | 1:500 | CST | 2955 |
| **GFP (4B10)** | Mouse | WB | 1:1000 | CST | 2955 |
| **GFP** | Mouse | WB | 1:4000 | Proteintech | 66002-1-Ig |
| **FLAG DYKDDDDK** | Mouse | IF | 1:500 | Proteintech | 66008-3-Ig |
| **FLAG DYKDDDDK** | Mouse | WB | 1:2000 | Proteintech | 66008-3-Ig |
| **FLAG DYKDDDDK (D6W5B)** | Rabbit | WB | 1:1000 | CST | 14793 |
| **Alexa Fluor 568 Anti-Rabbit (H+L)** | Goat | IF | 1:500 | Invitrogen | A11011 |
| **Alexa Fluor 488 anti-Rabbit IgG (H+L) SuperClonal** | Goat | IF | 1:500 | Invitrogen | A27034 |
| **Alexa Fluor 488 anti-mouse IgG (H+L)** | Rabbit | IF | 1:500 | Invitrogen | A27023 |
| **Alexa Fluor 647 anti-rabbit IgG (H+L)** | Goat | IF | 1:500 | Invitrogen | A21245 |
| **b-actin** | Mouse | WB | 1:1000 | Sigma | T7451 |
| **HA (C29F4)** | Rabbit | WB | 1:1000 | CST | 3724 |
| **HA** | Rabbit | WB | 1:5000 | Proteintech | 51064-2-AP |
| **HA** | Mouse | WB | 1:5000 | Proteintech | 66006-2-Ig |
| **His (27E8)** | Mouse | WB | 1:1000 | CST | 2366 |
| **6\*His** | Mouse | WB | 1:2000 | Proteintech | 66005-1-Ig |
| **HRP-conjugated GAPDH** | Mouse | WB | 1:4000 | Proteintech | HRP-60004 |

### Western Blotting

Cell lysates were subjected to SDS-PAGE on either self-cast 6, 8, or 13% polyacrylamide gels or pre-cast 4-20% Mini-PROTEAN^®^ TGX™ Precast Protein Gels (BioRad). Proteins were transferred to PDVF membranes by the wet transfer method at 50 V for 2 h. After transfer, membranes were washed for 5 min in TBST (Tris-buffered saline, 1% Tween-20), and blocked at room temperature for 1 h in TBST with 5% fat-free milk. After washing three times with TBST, membranes were incubated overnight at 4°C with primary antibodies, washed three times with TBST and incubated with the appropriate secondary HRP-conjugated anti-mouse or rabbit antibodies (BioRad) for 1 h at room temperature. Details of antibodies vendor and dilutions can be found in Supplementary Table 1. Membranes were then washed three times with TBST and developed using Clarity Western ECL Substrate (BioRad) on a ChemiDoc imaging system (BioRad) and using the Image Lab software (BioRad).

### FRET reduction after photobleaching (FRAP)

HEK 293 cells and *Ptch1^−/−^* MEFs were seeded in pre-coated, glass bottom, 35 mm, low, μ-Dishes (IBIDI) and incubated at 37°C, 5% CO_2_ until >90% confluent. Transient transfection of two different fluorescently-tagged PTCH1 and/or PTCH2-encoding plasmid was performed using Lipofectamine 2000 (HEK 293 cells) or Transit-X2 (*Ptch1^−/−^*MEFs). Approximately 24 h post-transfection, the cells were imaged using a Zeiss LSM 880 inverted confocal microscope at 37°C constant temperature. Cells were imaged using a pinhole of 1 airy unit (AU) and Plan-Apochromat 40x/1.4 Oil DIC objective. For detection of FRAP, GFP (donor) was excited at 488 nm and its emission intensity at 514 nm was measured before and after photo-bleaching of mCherry (acceptor). Donor and acceptor emission at 514 and 610 nm, respectively, were measured throughout to determine any increase in donor emission intensity post acceptor bleaching. Regions of interest (ROIs) were created to sample multiple bleaching and control events in individual image series acquisitions. Within each image series, a corresponding control (unbleached) ROI was obtained for each bleached sample ROI. Data was obtained over a series of 20 images taken at 1.2 sec intervals, 4 pre-bleach and 16 post-bleaching, to account for stage drift and observation of potential protein migration or recovery. All image series collected were saved as .czi files for subsequent processing across Zen Black (Zeiss), Excel 2016 (Microsoft) and Prism 8 (GraphPad). Fluorescence intensity values in the bleached ROI were normalised to the unbleached control ROI and expressed as a percentage. ROIs where mCherry bleaching was not achieved were discarded along with their control ROIs.

### FRAP Competition Assay

For FRAP competition assays, the two fluorescently-labelled plasmids that constitute the FRAP pair was co-transfected with a third plasmid encoding one of the PTCH variants lacking fluorescent protein tags or empty pcDNA3.1+ vector. The competing plasmid was co-transfected at a 4-fold mass excess. All microscope imaging and post-acquisition processing were as described before.

### Immunofluorescence

NIH 3T3 cells were seeded at 3 x 10^5^ cells /ml on 13 mm coverslips, in a 24 well plate containing DMEM, 10% BCS, 1% P/S. After 16 h, the cells were transfected with 1.5 μL of TransIT 2020 (Mirus) and 300 ng of plasmid DNA of interest in 50 μL of OptiMem per well. When the cells had reached confluence (∼24 h post-transfection) the media was changed to 0.5% BCS to induce ciliation. 48 h later the slides were washed with PBS (x3), fixed with 4% paraformaldehyde for 15 mins, followed by washing with PBS (x3) and permeabilisation with 0.2% Triton X-100 for 10 mins. After 3 x 5 min washes with PBS, the cells were blocked with 1% BSA in PBS for 1h at RT, followed by overnight incubation at 4 °C with anti-Arl13B antibody and one or more of the following primary antibodies: anti-myc tag clone 9B11, anti-GFP clone 4B10 or anti-FLAG, all diluted 1:500 in 1% BSA in PBS. The following day, the slides were washed with PBS 3 x 5 min and incubated with appropriate secondary antibodies (see Supplementary Table 1 for details) at 1:500 in 1% BSA in PBS, for 2 h in darkness. The slides were washed again carefully with PBS 3 x 5 mins and the coverslips were mounted onto glass microscope slides with Fluoroshield with DAPI (Sigma) and sealed with clear nail varnish. Confocal microscopy imaging was performed on a Zeiss LSM880 inverted confocal microscope using a 63x oil objective for visualisation of primary cilia and colocalisation with PTCH1 and PTCH2. Images were analysed using ImageJ with Fiji and Zeiss Blue software.

## RESULTS

### PTCH1 is found in homomeric state in live cells

To investigate whether the dimeric PTCH1 cryo-EM structures truly represent the quaternary structure of PTCH1 in live cells, we established a <u>F</u>örster resonance energy transfer (FRET) reduction <u>a</u>fter <u>p</u>hotobleaching (FRAP) system. Co-expression of PTCH1-mCherry and PTCH1-GFP, with the fluorescent proteins fused at the end of the C-termini allows to determine not only co-localisation in live cells, but also changes in FRET between mCherry and GFP if they are in very close proximity (<10 nm). Briefly, if the C-tails of PTCH1-mCherry and PTCH1-GFP are within 10 nm, GFP emission intensity is reduced by energy transfer to mCherry. Photobleaching of mCherry obliterates FRET, leading to increased GFP fluorescence (Fig. 1A). Time-lapsed detection of mCherry and GFP fluorescence intensity in live HEK 293 cells transfected with vectors encoding PTCH1-mCherry and PTCH1-GFP revealed a significant >10% net increase in GFP signal within 1.2 sec of mCherry photobleaching (Fig. 1B and C, green trace). The increase in GFP fluorescence intensity was not artefactual, since GFP fluorescence slowly decayed over time in the absence of photobleaching (Fig. 1B and C, grey trace). We reasoned that if the FRAP signal is due to physical interaction of PTCH1-mCherry and PTCH1-GFP and not to stochastic proximity of the monomers, the FRAP signal would be abolished by excess unlabelled protein. Therefore, we co-expressed PTCH1-mCherry and PTCH1-GFP with a 4-fold DNA mass excess of PTCH1-His or empty pcDNA3.1 vectors. FRAP was abolished by excess PTCH1-His but not by empty plasmid (Fig. 1D and E). Finally, we could also detect homomeric PTCH1 by co-immunoprecipitation (co-IP) of myc-tagged PTCH1 and HA-tagged PTCH1 (Fig. 1F). Altogether, these data indicate that PTCH1 exists in homomeric form (as a dimer or higher order oligomer) in the absence of Hh ligands in live cells.

**Figure 1.**
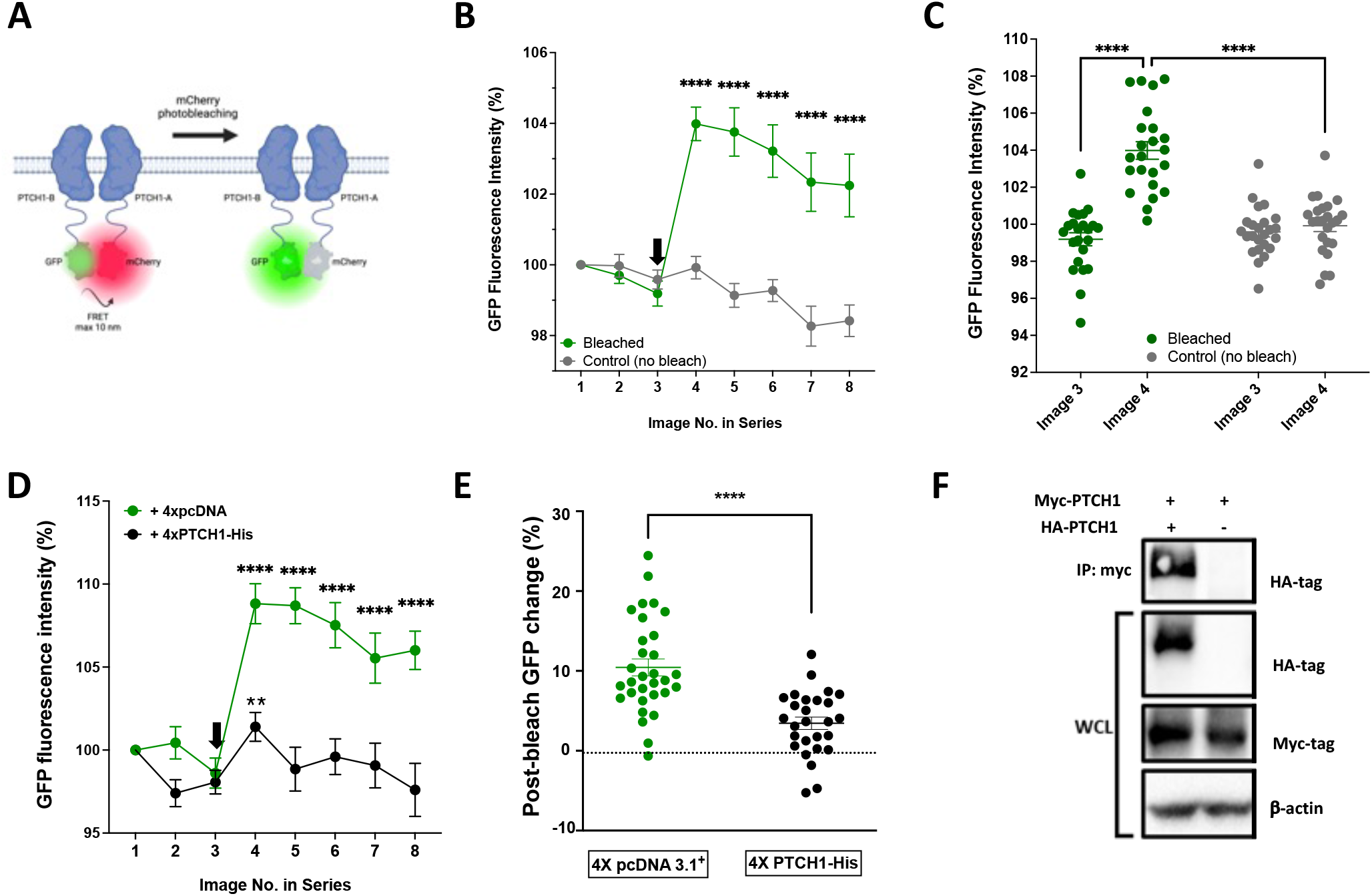
PTCH1 exhibits homomeric interactions in live cells. **A.** Schematic representation of the FRAP assay to detect direct protein-protein interactions within live cells. Two PTCH1 proteins, one containing a C-terminal mCherry and the other a C-terminal eGFP are co-expressed. If the two proteins directly interact, some emission energy from the eGFP is absorbed by mCherry. Photo-bleaching of mCherry prevents this excitation and the eGFP emission energy previously absorbed by mCherry can be detected as additional post-bleach eGFP emission intensity. Image created with BioRender.com. **B.** HEK 293 cells transiently co-expressing PTCH1-eGFP and PTCH1-mCherry, were subjected to confocal imaging with the FRAP interaction assay. Change in eGFP fluorescence intensity (expressed as % of initial fluorescence intensity) was measured across 8 acquisitions, with mCherry photo-bleached (green trace) or not bleached (grey trace) after the third image in the series. Statistical significance of indicated image number compared to image 3: **** P<0.0001 (n=23). **C.** Change in GFP fluorescence intensity between images 3 and 4 in HEK 293 cells in which mCherry was bleached or not (from Fig. 1B). **D.** Competition-based FRAP measurements in HEK 293 cells co-expressing PTCH1-eGFP and PTCH1-mCherry, when co-transfected with 4X mass excess of empty pcDNA 3.1^+^vector (green trace) or vector encoding PTCH1-His (black trace). Statistical significance of indicated image number compared to image 3 of the same group: ** P<0.01; **** P<0.0001 (n=27). **E.** Net change in GFP fluorescence intensity immediately post-bleach, corresponding to image 4 in the series, in PTCH1-eGFP and PTCH1-mCherry expressing HEK 293 cells with 4X excess of empty vector (green symbols) or 4X PTCH1-His (black symbols). **F**. Representative co-immunoprecipitation of HA-tagged PTCH1 with myc-tagged PTCH1 after co-expression in HEK 293 cells. WCL: whole cell lysate.

### Residues required for cholesterol transport are not involved in homomer formation

Since the sterol transport-like activity of PTCH1 could be necessary for homomer formation or stability, we systematically mutated residues predicted or shown to prevent PTCH1-dependent cholesterol redistribution and Gli transcriptional activity inhibition. X-ray crystallography studies of the homologous cholesterol transporter Niemann-Pick C1 (NPC1) revealed an exposed cavity within the SSD, predicted to accommodate a single cholesterol molecule (21). In agreement, mutations specific to the SSD of NPC1 have been found to prevent binding to a photoactivatable cholesterol analogue (22). We therefore hypothesised that key functional residues in the SSD of PTCH1 might be identifiable through homology modelling with NPC1. Threading the secondary structure of PTCH1 onto NPC1 (PDB ID code 5I31) using I-TASSER revealed mutational hotspots in both proteins within close proximity to the SSD cavity, which allowed identification of candidate residues involved in sterol transport (Fig. 2A). Proximity analysis identified several PTCH1 residues aligned to previously reported deleterious NPC1 mutations (Supplementary Table 2): A483, A484, T485, P490, L504 and F1109.

**Figure 2.**
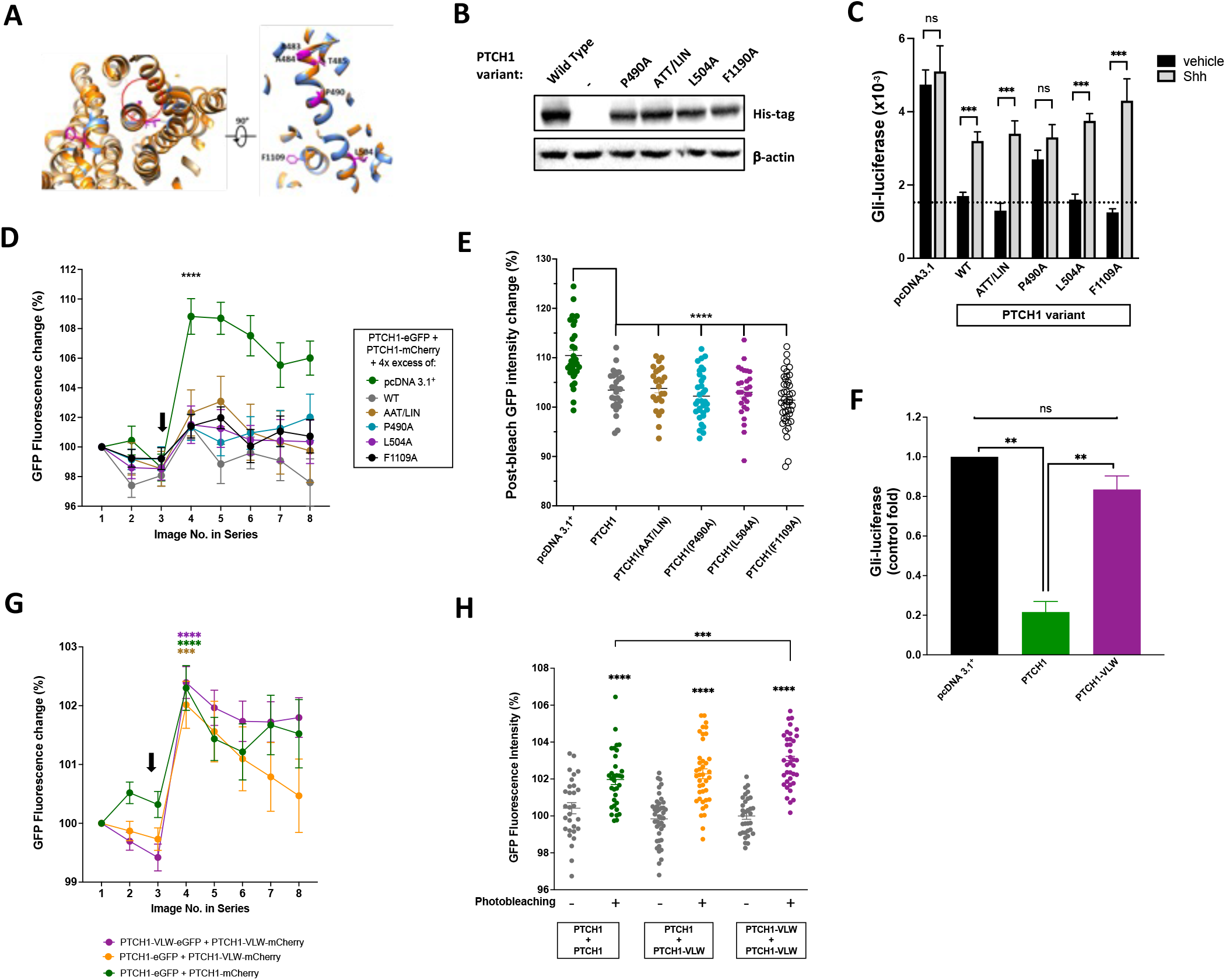
PTCH1 homomeric interaction is independent of activity. **A.** Alignment of NPC1 (PDB ID: 5U73) shown in orange, against the secondary sequence of PTCH1, threaded through NPC1 (PDB ID: 5U73) shown in blue. Residues selected for mutational investigation are shown with side chains in magenta. Left: bottom-up view of protein alignments, with the hydrophobic pocket of NPC1 indicated by a red circle. Right: restricted side view, depicting residues selected for mutational investigation. **B.** Western blot confirming comparable expression of wild type PTCH1 and the following mutants: P490A, AAT/LIN, L504A, F1109A in HEK 293 cells. **C.** Relative Gli-luciferase activity (Firefly/Renilla) in *Ptch1^−/−^* MEFs, co-transfected with empty vector (pcDNA3.1), PTCH1 or the indicated single or triple mutants in the absence (black) or the presence of Shh (grey). ns, not significant; *** P<0.001 (n=4). **D.** Competition-based FRAP of homomeric PTCH1-eGFP/PTCH1-mCherry interaction with 4X mass excess empty plasmid (pcDNA3.1), WT PTCH1 or the indicated mutants. **E.** Scatter graph of post-bleach GFP emission intensity, corresponding to image 4 in Fig. 2D. ****P<0.0001, n=28-43. **F.** Relative Gli-luciferase activity in *Ptc1^−/−^* MEFs, co-transfected with empty vector (pcDNA3.1), wild type PTCH1 or PTCH1-VLW mutant (n=3 biological repeats). ns, not significant; ** P<0.001. **G.** HEK 293 cells were transiently transfected with PTCH1-eGFP and PTCH1-mCherry (green trace), PTCH1-eGFP and PTCH1-VLW -mCherry (yellow trace) or PTCH1-VLW-eGFP and PTCH1-VLW-mCherry (purple trace) were subjected to the FRAP-based interaction assay. Change in eGFP fluorescence intensity (expressed as % of initial fluorescence intensity) was measured across 8 acquisitions, with mCherry photo-bleached after the third image in the series. Statistical significance of indicated image number compared to image 3: ***P<0.001; **** P<0.0001 (n=30-41). **H.** Change in GFP fluorescence intensity with the indicated FRET-pairs photobleached or not (related to Fig. 2G). ***P<0.001; ****P<0.0001 (n=30-41).

**Supplementary Table 2.** PTCH1 residues selected for mutagenesis in this study, based on known function, or phenotype given by mutations, on aligned residues in NPC1 and AcrB, identified by structural alignment.

| PTCH1 residue of interest | Known PTCH1 mutation? | Corresponding NPC1 residue | Known NPC1 mutation? | Corresponding AcrB residue |
| --- | --- | --- | --- | --- |
| A483 | A483T | L684 | L684F – Dystonia | T392 |
| A484 | - | I685 | I685T – Niemann-Pick disease C1 | L393 |
| T485 | T485I | V686 | V686M | T394 |
| P490 | P490Q | P691 | P691L - Niemann-Pick disease C1 | M398 (part of active site) |
| L504 | - | L705 | - | V412 |
| F1109 | - | G1195 | - | L972 (part of active site) |

We performed conservative mutations of those residues to avoid a large impact on TM core structure and protein stability on a PTCH1-K1426R background, which was originally generated for expression, purification, and structural work. The K1426R mutation increases PTCH1 plasma membrane retention and stability due to removal of a ubiquitylation site (20). Three single amino acid mutants (P490A, L504A and F1109A), and a triple mutant A483L, A484I, T485N (ATT/LIN) were generated. Western blotting confirmed comparable expression of all mutants (Fig. 2B). Only one of the mutants, PTCH1(P490A), showed impaired activity in GLI-luciferase assays in *Ptch1^−/−^* MEFs (Fig. 2C). In addition to a reduced inhibitory activity, the mutant is insensitive to the presence of Shh, while the other mutants display normal activity and responsiveness to Shh (Fig. 2C). We next tested if the reduced activity mutant PTCH1(P490A) was able to displace FRAP between PTCH1-mCherry and PTCH1-GFP using the FRAP competition assay discussed earlier. As shown in Fig. 2D and E, a 4-fold mass excess of PTCH1(P490A) abolished the FRAP signal in a similar way than wild type PTCH1 and the additional three mutants, suggesting that the reduced activity of PTCH1(P490A) does not affect homomeric state formation in a cellular context.

Next, we generated a triple mutant that was reported to abolish PTCH1 activity based on perturbation of the sterol-binding pocket in the neck region of the ECD (named site II) (4). The PTCH1 mutant V111F, L114F and W115A (referred here as VLW) showed a complete loss of function in GLI-luciferase assays (Fig. 2F). Thus, we used this mutant to investigate if activity is required for formation of stable homomers. In the FRAP system described before, the increase in GFP fluorescence intensity between monomers of PTCH1(VLW) was statistically larger than that of monomers of wild type PTCH1, while combination of wild type and mutant PTCH1 had an intermediate FRAP signal (Fig. 2G and H). Altogether, these findings strongly indicate that PTCH1 exists in homomeric state in the absence of ligand, regardless of its activity. Furthermore, the results suggest a dynamic equilibrium in which the inactive state, which likely has structural similarities to the Shh-liganded PTCH1 asymmetric dimer, might have a higher propensity for self-interaction.

### PTCH2 also exists in homomeric form in live cells

We extended the FRAP system to study the quaternary structure of the understudied PTCH2 isoform. As with PTCH1, co-transfection of HEK 293 cells with PTCH2-mCherry and PTCH2-GFP allowed us to detect a FRAP signal using time-lapsed imaging (Fig. 3A and B). In agreement, co-IP studies showed stable physical interaction between HA- and FLAG-tagged PTCH2 variants (Fig. 3C). Sequence alignment and threading of PTCH2 onto PTCH1 identified L82, L85 and W86 as equivalent residues to the V111, L114 and W115 triad of PTCH1 essential for activity (Fig. 3D). We predicted that mutation of the equivalent residues in PTCH2, generating the PTCH2 L82F, L85F, W86A (LLW) mutant, would abolish the potential sterol transport activity of PTCH2 and, therefore, impair its activity to rescue GLI-luciferase activity in *Ptch1^−/−^* MEFs. However, in stark contrast with PTCH1 VLW, the PTCH2 LLW mutant did not impair PTCH2 activity in the Gli-luciferase assay, which in our hands is significantly lower than the activity of the PTCH1 isoform at saturating levels (Fig. 3E). As expected, based on the behavior of PTCH1, the PTCH2 (LLW) mutant did not prevent homomer formation with wildtype PTCH2 or with itself (Fig. 3F and G).

**Figure 3.**
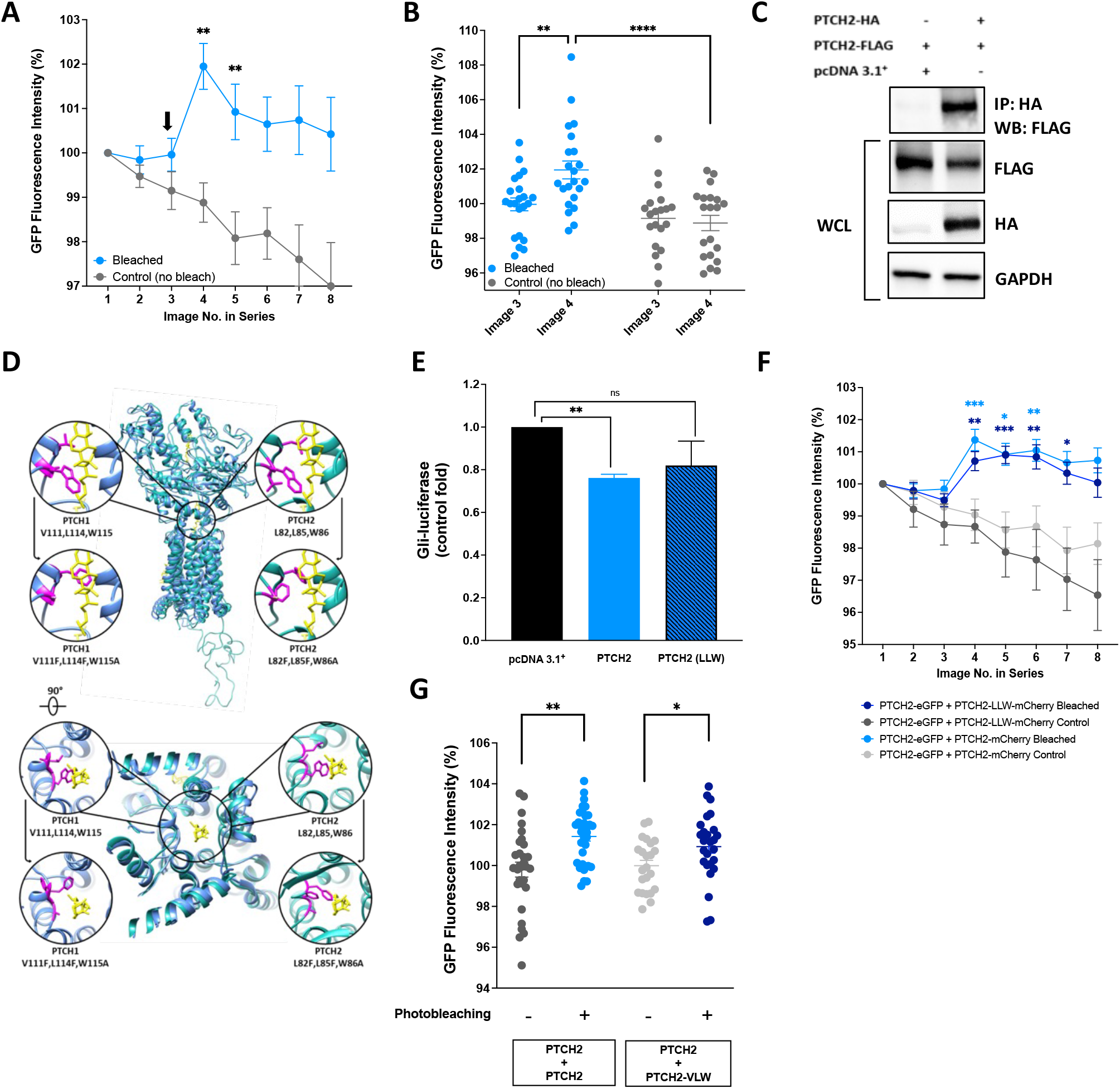
PTCH2 forms homomeric complexes in the active state. **A.** HEK 293 cells transiently expressing PTCH2-eGFP and PTCH2-mCherry, were subjected to confocal imaging with the FRAP interaction assay. Change in eGFP fluorescence intensity (expressed as % of initial fluorescence intensity) was measured across 8 acquisitions, with mCherry photo-bleached (blue trace) or not bleached (grey trace) after the third image in the series. Statistical significance of indicated image number compared to image 3 bleached: ** P<0.005 (n=22). **B.** Change in GFP fluorescence intensity between images 3 and 4 in HEK 293 cells in which mCherry was bleached or not (from Fig. 3A) **P<0.005; ****P<0.0001 (n=22). **C.** Representative co-immunoprecipitation of FLAG-tagged PTCH2 with HA-tagged PTCH2 after co-expression in HEK 293 cells. WCL: whole cell lysate. **D.** Side and top view of the 3D structure of PTCH1 (PDB ID: 6MG8) shown in blue, with the threaded PTCH2 (UniProtKB: Q9Y6C5) sequence in turquoise. The putative sterol channel is indicated by a black ring, with zoom views of the hydrophobic tunnel residues in PTCH1 (top left) and predicted equivalent residues in PTCH2 (top right), in magenta. Cholesterol-like densities are displayed in yellow. The top zoom windows display original residues and their side chains, whilst the windows beneath display the predicted mutational alterations to side chains. Structural analysis and alterations were performed in Chimera (23). **E.** Relative Gli-luciferase activity in *Ptch1^−/−^* MEFs co-transfected with empty vector (pcDNA3.1), wild type PTCH2 or PTCH2-LLW mutant (n=3 biological repeats). ns, not significant; *** P<0.005. **F.** HEK 293 cells were transiently transfected with PTCH2-eGFP and PTCH2-mCherry (blue trace) or PTCH2-eGFP and PTCH2-LLW-mCherry (dark blue trace) were subjected to the FRAP-based interaction assay. Change in eGFP fluorescence intensity (expressed as % of initial fluorescence intensity) was measured across 8 acquisitions, with mCherry photo-bleached after the third image in the series. Grey traces show change in GFP fluorescence in unbleached adjacent ROIs. Statistical significance of indicated image number compared to bleached image 3: *P<0.05; **P<0.005; *** P<0.0001 (n=24-32). **G.** Change in GFP fluorescence intensity in the indicated FRET pairs photobleached or not (from Fig. 3F), * P<0.05; ** P<0.005.

### PTCH1 and PTCH2 form active heteromeric complexes in live cells

Given the capacity of both PTCH1 and PTCH2 to form homomers, we investigated the potential of heteromeric complexes and their properties. We first tested the interaction between PTCH1 and PTCH2 by co-IP. Epitope-tagged fusions of PTCH1 (myc-PTCH1) and PTCH2 (FLAG-PTCH2) were co-transfected into HEK 293 cells or co-transfected individually with empty vector. PTCH1 pulldown immunoprecipitated PTCH2, while the reverse co-IP confirmed that PTCH2 was able to pulldown PTCH1 (Fig. 4A). Interestingly, PTCH2 steady state levels were reduced when co-expressed with PTCH1, suggesting that interaction with PTCH1, which has a short half-life controlled by two PPXY motifs that interact with ITCH and other ubiquitin E3 ligases, reduces PTCH2 stability.

**Figure 4.**
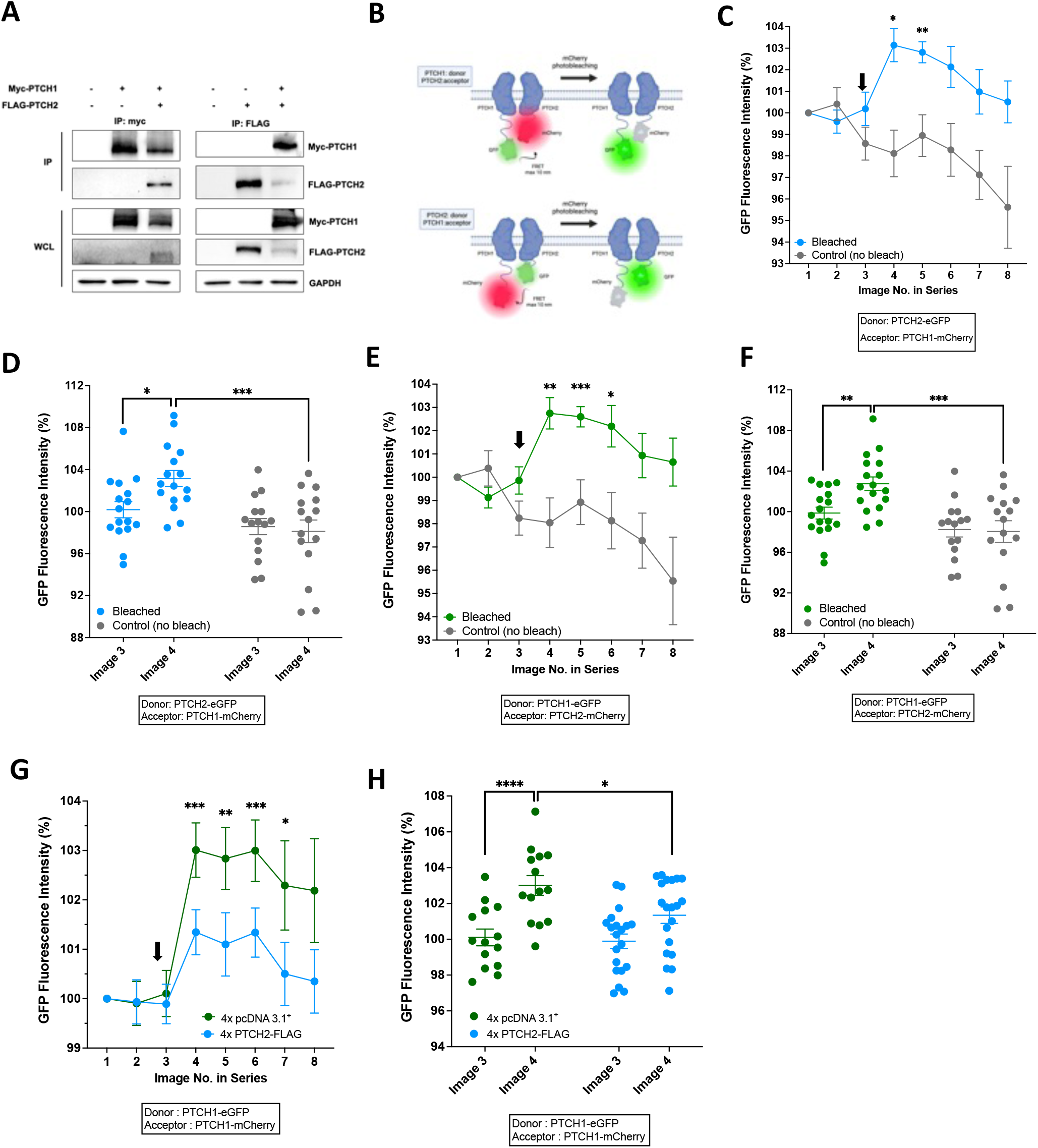
PTCH1 and PTCH2 form heteromeric complexes in live cells. **A.** Two-way co-immunoprecipitation of myc-PTCH1 and FLAG-PTCH2 after co-expression in HEK 293 cells. IP: immunoprecipitation; WCL: whole cell lysate. **B.** Schematic representation of the FRAP assay to detect interaction between PTCH1 and PTCH2, using either of the proteins as donors or acceptors, created with BioRender.com. **C.** Change in GFP fluorescence intensity in HEK 293 cells co-expression PTCH2-eGFP and PTCH1-mCherry upon photobleaching of mCherry after the third image (blue trace) or in control unbleached regions (grey trace). * P<0.05; ** P<0.01 (n=15-16). **D.** Change in GFP fluorescence intensity between image 3 and 4 in regions subjected to photobleaching (blue symbols) or not (grey symbols) (from Fig. 4C), * P<0.05; ** P<0.001 (n=15-16). **E.** Change in GFP fluorescence intensity in HEK 293 cells co-expressing PTCH1-eGFP and PTCH2-mCherry upon photobleaching of mCherry after the third image (green trace) or in control unbleached regions (grey trace). ** P<0.005; *** P<0.001 (n=15-17). **F.** Change in GFP fluorescence intensity between image 3 and 4 in regions subjected to photobleaching (green symbols) or not (grey symbols) (from Fig. 4E), ** P<0.005; *** P<0.001 (n=15-17). **G.** Competition-based FRAP of homomeric PTCH1-eGFP/PTCH1-mCherry interaction with 4X mass excess empty plasmid (pcDNA3.1; green trace) or PTCH2-FLAG (blue trace). * P<0.05; ** P<0.005; ***P<0.0005 (n=14-20). **H.** Change in GFP fluorescence intensity in HEK 293 cells co-expressing PTCH1-eGFP and PTCH1-mCherry upon photobleaching of mCherry after the third image with a 4X excess of empty vector (pcDNA3.1, green symbols) or a 4X excess of PTCH2-FLAG (blue symbols). * P<0.05; *** P<0.001 (n=15-17).

PTCH1 and PTCH2 heteromeric interactions were further investigated using the FRAP assay. To exclude any possible introduction of bias, through non-specific effects of unequal expression levels and basal fluorescence intensity, both PTCH1 and PTCH2 were tested in the role of donor (eGFP) and acceptor (mCherry) (Fig. 4B). Photobleaching of mCherry resulted in a significant increase in normalised GFP fluorescence intensity in both co-transfected conditions; PTCH1-eGFP + PTCH2-mCherry, and PTCH2-eGFP + PTCH1-mCherry (Fig. 4 C-F). The magnitude of GFP signal change was ∼50% lower of that of PTCH1 homodimers, which might reflect the increased distance of the fluorescent proteins given that the C-tail of PTCH1 is 273 residues while the C-tail of PTCH2 is 23 amino acids long (Fig. 4B). Finally, we tested whether excess unlabelled PTCH2 was able to disrupt PTCH1 homomers. Co-transfection of 4-fold excess FLAG-PTCH2 significantly reduced the FRAP signal between PTCH1-eGFP and PTCH1-mCherry, unlike a 4-fold excess of empty vector (Fig. 4G and H), suggesting that the relative abundance of PTCH1 and PTCH2 controls the ratio of homo to heteromeric complexes.

### Heteromeric complexes of PTCH1 and PTCH2 display synergistic activity

Given the significant differences in GLI-inhibitory activity of PTCH1 and PTCH2, we sought to investigate the activity of PTCH1-PTCH2 heteromers compared to each isoform alone. For these experiments, only 50% of the plasmids encoding PTCH1 or PTCH2 were transfected along the same amount of empty vector into *Ptch1^−/−^* MEFs. Transfection of 50% amount of the PTCH1 plasmid was used to achieve sub-maximal inhibitory activity (Fig. 5A). Instead, transfection of 50% of PTCH2 plasmid was insufficient to allow detection of statistically significant activity, given its much lower maximal activity (Fig. 5A). Surprisingly, co-transfection of 50% PTCH1 and 50% PTCH2 resulted in greater Gli-luciferase inhibitory activity than 50% of PTCH1 alone (Fig 5A, blue bar). Under the assumption that heteromeric complexes can be represented by asymmetric dimers, as observed by cryo-EM of purified PTCH1 in complex with Shh, and that formation of homo and heterodimers has the same probability, cells co-expressing both isoforms should contain 25% PTCH1 homodimers, 25% PTCH2 homodimers and 50% PTCH1-PTCH2 heterodimers. If heteromeric complexes had lower activity than PTCH1 homodimers, we would expect a “dilution” of PTCH1’s activity, and higher GLI-luciferase signal when PTCH1 and PTCH2 are co-expressed compared to PTCH1 alone. The potentiation of GLI-luciferase inhibition suggests that the heteromeric PTCH1-PTCH2 have higher intrinsic activity than each isoform in homomeric form.

**Figure 5.**
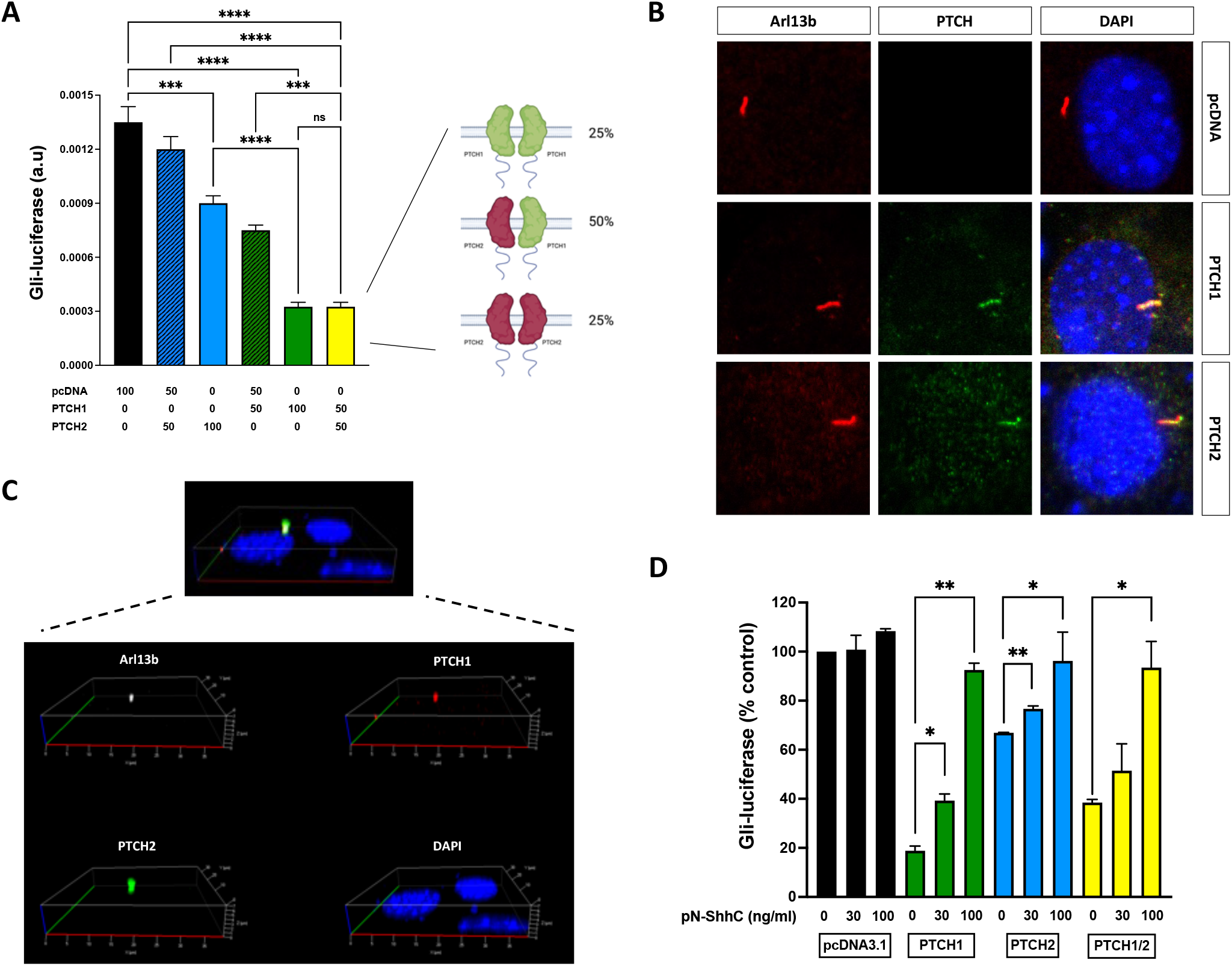
Heteromeric PTCH1-PTCH2 complexes co-localise to cilia and display high activity. **A.** Relative Gli-luciferase activity in *Ptch1^−/−^* MEFs, co-transfected with the indicated percent amounts of empty vector (pcDNA3.1), PTCH1, or PTCH2, alone or in combination. Ns, not significant; *** P<0.0005; ****P<0.0001 (n=3 independent experiments). Schematic created with BioRender.com. **B.** Ciliary localisation of PTCH1 and PTCH2. NIH 3T3 cells were transfected with PTCH1-eGFP or PTCH2-eGFP, serum starved, and 24 h later stained with Arl13b (ciliary marker) or anti-GFP. Nuclei are counterstained with DAPI. **C.** Co-localisation of PTCH1 and PTCH2 in primary cilia. NIH 3T3 cells were co-transfected with PTCH1-eGFP and PTCH2-FLAG and serum starved to induce ciliogenesis. After 24 h, cells were stained with Arl13b (ciliary marker), anti-GFP (PTCH1) and anti-FLAG (PTCH2). The image shows a Z-stack of a representative cilium co-expressing the three markers. **D.** Induction of Gli-luciferase activity by mid- and high concentrations of pN-ShhC in *Ptch1*^−/−^ MEFs transiently transfected with PTCH1, PTCH2 or a 50% PTCH1 plus 50% PTCH2. * P<0.05; **P<0.01 (n=3 independent experiments).

This prompted us to investigate if the presence of PTCH1 is necessary for primary cilia localisation of PTCH2, which could explain the increased activity of PTCH2 as part of heterodimers compared to homomeric PTCH2. Thus, we investigated cilia localisation of PTCH1 and PTCH2 in serum-starved NIH 3T3 cells using the primary cilia marker Arl13b. As shown in Fig. 5B, when individually expressed in NIH 3T3 cells, both PTCH1 and PTCH2 are readily detected in primary cilia. Similar findings were observed in HEK 293 cells and *Ptch1^−/−^*MEFs (Suppl. Fig. 1). This result indicates, first, that the lower activity of PTCH2 cannot be ascribed to inability to traffic to the right compartment and, second, that PTCH2 does not require heterodimerisation with PTCH1 to accumulate at the primary cilium. In addition, co-expressed PTCH1 and PTCH2 co-localised with Arl13b in primary cilia (Fig. 5C). Z-stack projections showed a similar distribution of PTCH1 and PTCH2 along the length of the ciliary axoneme (Fig. 5C). Altogether, the data support a model of pre-existing PTCH1/PTCH2 heterodimers in the primary cilium with high Smoothened repressive activity in the absence of Hh ligands.

Previous reports of similar affinity of Shh to PTCH1 or PTCH2 used competition of binding of either non-lipidated ^125^I-Shh (24) or non-lipidated Shh-Fc fusions (25). However, the current understanding of the asymmetric mode of binding of dually-lipidated Shh (pNShhChol) to the PTCH1 homodimer provided by cryo-EM strongly suggests that the binding affinity assays done in the past would reflect only the affinity of the globular domain of Shh ligands to the Patched-B subunit of homo- or heterodimers of different composition, since neither of the ligand variants used would be able to engage the Patched-A subunit by the pincer grasp mechanism (8). Therefore, we revisited the potency of the physiological pNShhC ligand to inhibit heterodimers and the two types of homodimers. The concentrations of pNShhC used were selected based on their activity in GLI-luciferase activity assays in NIH 3T3 cells: 30 ng/mL increased Gli-luciferase by ∼50% of maximum representing the EC_50_, and 100 ng/mL proved to be a saturating concentration. Our results indicate that PTCH1-PTCH2 heteromeric complexes have a similar response to lipidated Shh than PTCH1 or PTCH2 individually, reaching the same level of maximal GLI-luciferase activity (Fig. 5D). Therefore, formation PTCH1-PTCH2 heteromers might reduce the functional consequence of deleterious mutations in either of the isoforms by displaying strong SMO inhibitory activity while maintaining a physiological responsiveness to SHH ligand.

### Sterol transport activity of PTCH1, but not PTCH2, is essential for heterodimers activity

We previously showed that the hydrophobic tunnel mutants of PTCH1 and PTCH2 (PTCH1(VLW) and PTCH2 (LLW)) have very low activity in the GLI-luciferase assay (Fig. 2F and 3E). However, the low activity of wildtype PTCH2 and the similarity with the PTCH2 (LLW) mutant suggests that, perhaps, PTCH2 has a much smaller or null cholesterol transport activity than PTCH1. This could explain why PTCH2 is not redundant with PTCH1 in knockout genetic mouse models. To further investigate this possibility, we measured total cholesterol changes in HEK 293 cells transfected with a control plasmid, PTCH1, PTCH2 and the two mutants of the hydrophobic tunnel by filipin staining. Overexpression of PTCH1 significantly increased filipin staining compared to empty vector (Fig. 6A). The PTCH1 (VLW) mutant, as expected, did not show cholesterol accumulation like wildtype PTCH1, demonstrating the validity of this approach as surrogate measure of PTCH cholesterol transport activity. In agreement with the lack of effect of the triple LLW mutation in PTCH2, both wildtype and mutant PTCH2 show negligible accumulation of cholesterol in comparison to PTCH1 (Fig. 6A). These data suggest that only PTCH1 can function as a cholesterol transporter, or that the sterol transport-like activity of PTCH1 is much higher than that of PTCH2.

**Figure 6.**
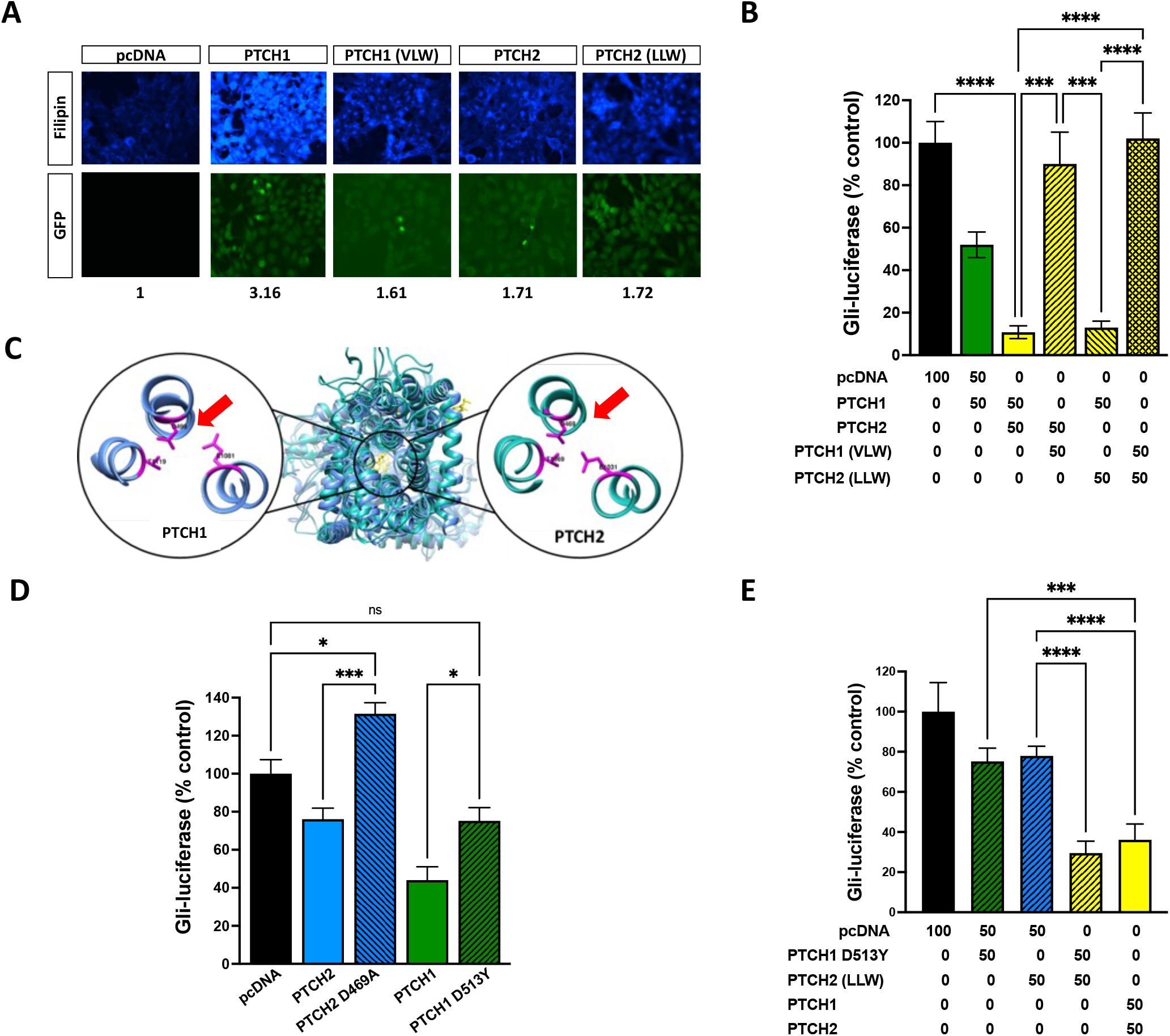
Cholesterol and cation transport requirements for activity of homomeric and heteromeric complexes. **A**. Free cholesterol staining with filipin in HEK 293 cells transfected with empty vector (pcDNA) or with GFP-tagged constructs of wild type PTCH1, PTCH1 (VLW), wild type PTCH2 or PTCH2 (LLW). Numbers indicate filipin intensity compared to pcDNA control. **B**. Gli-luciferase activity of the same PTCH variants alone and in combination transfected in the indicated percent amounts into *Ptch1*^−/−^ MEFs. *** P<0.0005; ****P<0.0001 (n=3-5 independent experiments). **C.** Comparison of the predicted position of PTCH2-D469 to that of PTCH1-D499. The Aspartic acid in question and two known interaction partners are shown with side chains in magenta. PTCH1 (PDB ID: 6MG8) is shown in blue. PTCH2, shown in turquoise, represents the secondary structure, threaded into the template structure of PTCH1. The protein structures are depicted from a bottom view. **D.** Gli-luciferase activity of PTCH2, PTCH2 D469A, PTCH1 and PTCH1 D513Y in *Ptch1*^−/−^ MEFs. Ns, not significant; * P<0.05; *** P<0.0005 (n=3 independent experiments). **E.** Gli-luciferase activity of the indicated PTCH1 and PTCH2 variants transfected alone and in combination in the indicated percent amounts into *Ptch1*^−/−^ MEFs. *** P=0.0001; ****P<0.0001 (n=4 independent experiments).

Based on these observations, we hypothesised that the activity of PTCH1-PTCH2 heteromeric complex relies on the sterol transport-like function of the PTCH1 monomer. Thus, we compared the GLI-inhibitory activity of the wildtype heterodimers to that of heterodimers in which one of the partners cannot transport cholesterol. As predicted, co-expression of wildtype PTCH2 and PTCH1 (VLW) resulted in formation of inactive heterodimers, while PTCH2 (LLW) mutant was able to synergise with wildtype PTCH1 just like wildtype PTCH2 (Fig. 6B). This data indicates that the functionality of the PTCH1-PTCH2 heterodimer strictly requires the cholesterol transport activity of the PTCH1 subunit.

Since PTCH proteins are evolutionary related to bacterial RND permeases, which use energy stored in ionic gradients to mobilise hydrophobic compounds, and the conserved GxxxDD motif that lines the cation channel is essential for PTCH1 activity (4, 18), we investigated if it is also necessary in PTCH2. Sequence analysis and threading of PTCH2 onto PTCH1 revealed conservation of the GxxxDD motif in the same location (Fig. 6C). The “cation gate” mutant PTCH2 D469A lacks GLI-inhibitory activity and apparently is dominant negative over endogenous Ptch2 in *Ptch1^−/−^* MEFs, since GLI-luciferase is consistently ∼30% higher in the presence of PTCH2 D469A compared to empty vector (Fig. 6D). Conversely, the equivalent PTCH1 D513Y mutant shows reduced but not full loss of activity (Fig. 6D). Given that the PTCH2 (LLW) mutant can form active heterodimers with wildtype PTCH1, we hypothesised that heterodimers might provide robustness to HH signalling regulation by PTCH by providing cholesterol and cation transport in *trans*; i.e. with each monomer providing one of the activities. We tested this hypothesis by co-expressing PTCH1 D513Y and PTCH2 (LLW), which individually display very low activity in *Ptch1^−/−^* MEFs (Fig. 6E). The two inactive proteins were able to reconstitute the SMO inhibitory activity to the same level than the wild type monomers, as seen by reduced GLI-luciferase activity (Fig. 6E), suggesting a coupling mechanism by which cation transport by PTCH2 provides the free energy for cholesterol mobilisation by PTCH1. This model would require the existence of stable heterodimers in the absence of Hh ligand, as suggested by our FRAP and co-IP studies. In summary, these results support the notion that the activity of heterodimers relies exclusively on PTCH1 cholesterol transport activity, which can be powered by ionic coupling by either of the two monomers.

## DISCUSSION

In the recent years, several groups reported the near atomic structure of purified complexes of the main HH ligand receptor in vertebrates PTCH1 bound to its inhibitory ligand Shh using cryo-EM (7, 8). The inhibited structure revealed two monomers of PTCH1 orientated asymmetrically between themselves bridged by dually lipidated Shh bound to the PTCH1-A monomer by the globular calcium-mediated interphase and to PTCH1-B by an extended arm formed by the most N-terminal residues and the N-terminal palmitate. The purified Shh-bound dimer was shown to form a loosely associated “dimer or dimers” in which 2 PTCH1:1 Shh complexes interact with another 2 PTCH1:1 Shh complex through the extracellular domain 2 (ECD2) (7). The inhibited structure immediately suggests that cholesterol transport is likely to occur asymmetrically as well and raised the question of the quaternary structure of PTCH1 in its active state. Here, we show that active PTCH1 exists as a homodimer or higher order oligomers (possibly a tetramer) using a combination of co-immunoprecipitation, FRET-recovery after photobleaching in live cells, and functional studies in the absence of a bridging ligand. We also demonstrated that the less studied isoform PTCH2, which is dispensable for embryonic development but was shown to be an important positive modifier of PTCH1 activity in a *Ptch1* heterozygote or null background (15, 17), also forms homomeric complexes with Smo-regulatory activity in live cells. Unlike PTCH1, PTCH2 overexpression has minimal impact on free cholesterol levels and mutation of residues predicted to surround a sterol cavity do not affect its activity. Free cholesterol, or chemically-active cholesterol, is unesterified and uncomplexed to sphingomyelin, and can be detected using filipin, a microbial derived macrolide. Previous elegant studies have shown that PTCH1 reduces free “accessible” cholesterol in the outer lamella of the plasma membrane, preventing cholesterol-dependent activation of Smo (12). Mutation of V111, L114 and W115 in PTCH1, which form a cavity surrounding a sterol-like density in the cryo-EM structure (4) reduces PTCH1 function to inhibit Gli-luciferase activity and reduces filipin staining in cells, suggesting that it impairs cholesterol transport by PTCH1.

Early conservation and homology studies showed that PTCH proteins are related to RND bacterial permeases, which transport hydrophobic small molecules using a coupled proton-motive force. Proton transport requires a conserved hydrophilic pore composed of a GxxxDD motif, which we refer here as the “cation gate” (26). The GxxxDD motif is conserved in PTCH1 and PTCH2, and mutation of any of the triad residues in PTCH1 has been shown to abolish its activity (4, 18). In agreement, a PTCH1 D513Y missense mutation described in individuals affected by Gorlin syndrome (19), displayed significantly less activity than wild type PTCH1 in our assays, but not null. Interestingly, despite the apparent low cholesterol transport activity of PTCH2, we found that the D469A mutation (predicted to be in the same position as D513 of PTCH1) creates a protein with dominant negative behavior, suggesting an important role of PTCH2 as cation transporter. The PTCH2 D469A mutant had been previously reported to exert dominant negative effects in *Ptch1^−/−^*MEFs and was proposed to regulate PTCH1 activity *in trans* (18). In this study, we demonstrate for the first time that PTCH1 and PTCH2 form active heteromeric complexes that depend on the cholesterol transport activity of PTCH1 but in which cation transport can be provided by either PTCH1 or PTCH2. The latter can explain how PTCH2 can regulate PTCH1 in *trans*, by providing additional cation transport that powers PTCH1 sterol transport. However, this model requires significant interaction between the PTCH isoforms, or molecular crowding, to allow for antiporter transport to occur through two separate molecules. Our finding that a cation transport-impaired PTCH1 mutant (D513Y) and a cholesterol transport-impaired PTCH2 mutant (LLW) can reconstitute active Smo/Gli inhibitory activity support this working model (Fig. 7).

**Figure 7.**
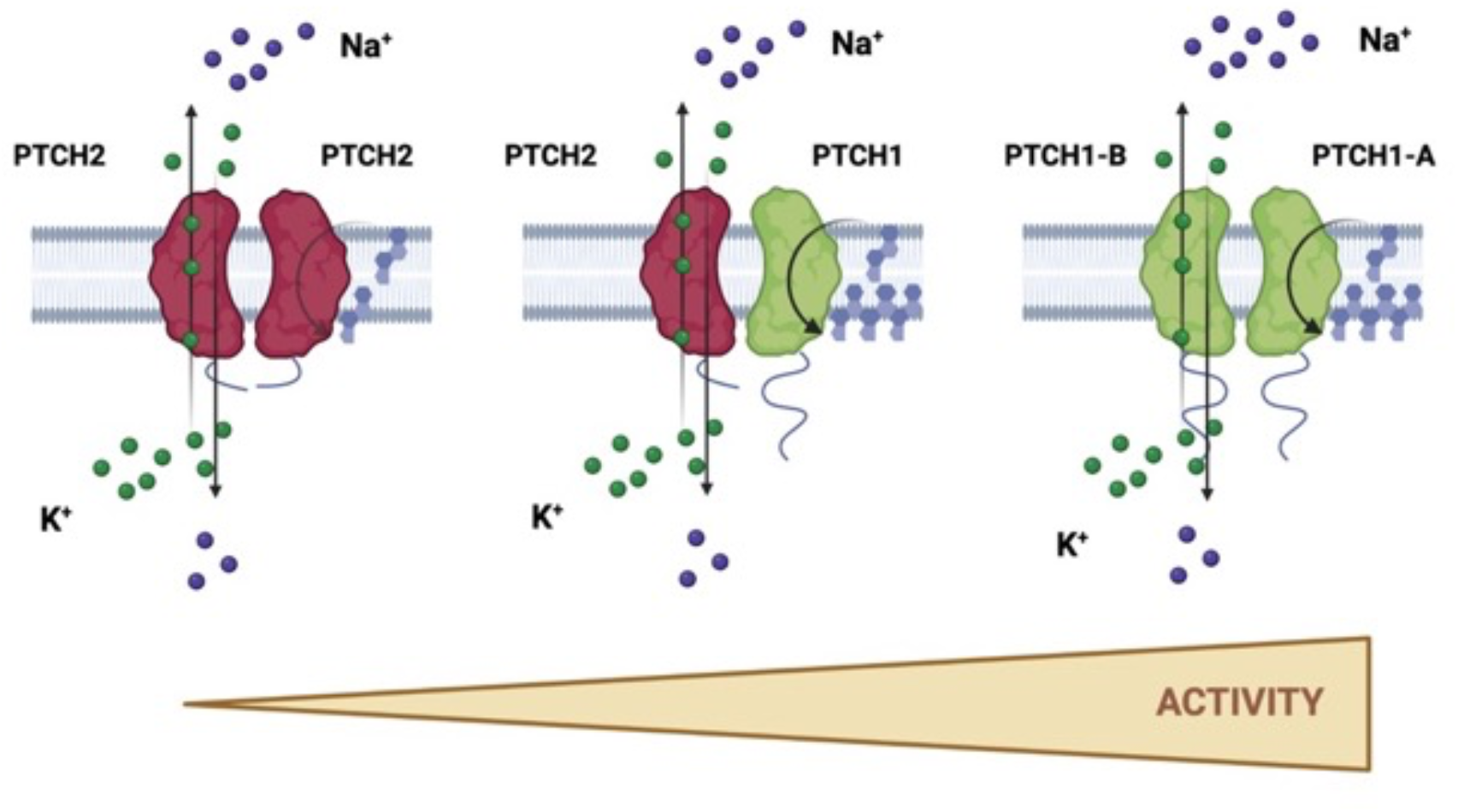
Proposed hypothetical conformation of the active PTCH1 and PTCH2 homomeric and heteromeric complexes. PTCH1 and PTCH2 co-exist as homo or heterodimers (or higher order complexes) in which each monomer adopts an asymmetric function, the monomer equivalent to PTCH1-B in the inactive conformation bound to Shh is posed for cation transport (K^+^ or Na^+^) and the monomer equivalent to PTCH1-A exerts the cholesterol transport activity. PTCH2 homodimers have very low cholesterol transport activity, which is translated as low Smo inhibitory capacity, while both heterodimers and PTCH1 homodimers have maximal activity. Created with BioRender.com.

Previous work proposed that *Drosophila* Patched could be a trimer, brought together by physical interaction of the C-terminal domains (27). Most of the related bacterial RND permeases are trimers, although some are dimers or even monomers (28); however, our assays do not permit distinguishing the stoichiometry of the homo and heteromeric complexes. Our unpublished data indicates that the physical interaction in both homomeric and heteromeric complexes does not require the large intracellular loop and the C-tail (A.J.T and N.A.R.-D.G., manuscript in preparation), suggesting that either the 12-TM core and/or the extracellular domains, as suggested by the cryo-EM tetramer, play a key role in facilitating complex formation. We believe it is more likely that PTCH exists as pre-formed dimers, given the structure of the inhibited conformation with Shh, but cannot rule out a larger reorganisation. Transporters typically cycle between active and inactive conformations. In support, using a nanobody that stabilises the active PTCH1 conformation, it was proposed that a helix switch in ECD2 proximal to the lipid membrane distinguishes between the active and inactive conformations (29). We believe it is likely that a dimeric complex cycles between the active and inactive conformation and that the latter is stabilised by binding of a Hh ligand.

The existence of active PTCH1-PTCH2 complexes might provide a backup way of reducing the impact of some deleterious mutations in PTCH1, but not those that impair cholesterol transport. The first evidence of a physical interaction between PTCH1 and PTCH2 was reported by co-immunoprecipitation (30), one of the multiple methods used in this study to demonstrate the existence of heteromers. Our findings also suggest that the expression level of PTCH2 can play an important regulatory role. While PTCH1 is essential for embryonic development and has a well-characterised tumour suppressor function in adult tissues, PTCH2 is mostly dispensable when PTCH1 is present. The much lower activity of PTCH2 supports this notion of dispensability. Nonetheless, mutations of PTCH2 were linked to rare cases of Gorlin syndrome, highlighting that even in the presence of wild type PTCH1, PTCH2 plays an important regulatory role. In one study, an autosomal dominant mutation PTCH2 (R719Q), a highly conserved residue in the ECD2 which is also present in PTCH1, was the only Hh pathway alteration in a family with mild Gorlin syndrome symptoms (31). A latter case report identified a healthy female with a homozygous PTCH2 frameshift-causing mutation (32); however, it was not explored if the frameshift led to loss of PTCH2 expression as it could trigger differential splicing. The fact that a point mutation in PTCH2 can dysregulate Hh signalling while complete loss of PTCH2 does not, supports our findings of existence of active heteromeric complexes. Moreover, it is possible that some mutations in PTCH1 are functionally silent by a built-in redundancy provided by interaction with PTCH2. The active heteromeric model can explain the large increase in medulloblastoma, sarcoma and basal cell carcinoma tumour burden, and reduced latency of appearance, in *Ptch1^+/−^* mice when an allele of *Ptch2* is inactivated (15). Importantly, the tumours in *Ptch1^+/−^Ptch2^+/−^* mice are very similar to those in *Ptch1^+/−^* animals, suggesting that Ptch2 is a positive modulator of Ptch1 function. Ptch2 also plays a key role in the absence of Ptch1, severely affecting digit patterning and reducing the outgrowth of the forelimb bud (17).

Both homo and heteromeric complexes can be inhibited by dually-lipidated Shh with similar potency. Because the heteromeric complex depends on PTCH1’s cholesterol transport capacity, we speculate that the palmitate-bearing arm of Shh which is essential and sufficient to stimulate canonical Hh signalling, will always interact with PTCH1 while the globular Ca^2+^-binding interphase will interact with PTCH2. Future structural biology work will provide essential insights into the mechanism of PTCH1-PTCH2 heterodimer function and will inform unknown functional asymmetries in the PTCH1 dimer.

## ACKNOWLEDGEMENTS

This research was funded by the Biotechnology and Biological Sciences Research Council, grant number BB/S01716X/1 to N.R.D-G. and D.S.G., a University of Leeds Scholarship to A.J.T., and a School of Molecular and Cellular Biology scholarship to F.C. We are indebted to Dr. Ruth Hughes from the Faculty of Biological Sciences Bioimaging facility for support with the Zeiss LSM880 Airyscan confocal microscope, funded by a Wellcome Trust grant (WT104918MA).

## AUTHORS CONTRIBUTION

Conceptualization, N.A.R.-D.G.; methodology, N.A.R.-D.G., A.J.T., and C.A.J.; investigation, A.J.T., F.C., D.S.G. and H.O.; data analysis, N.A.R.-D.G., A.J.T., F.C., D.S.G. and H.O.; writing—original draft preparation, A.J.T. and N.A.R.-D.G.; writing—review and editing, A.J.T. and N.A.R.-D.G.; visualization, A.J.T., F.C., D.S.G. and H.O.; supervision, N.A.R.-D.G. and C.A.J.; funding acquisition, N.A.R.-D.G. All authors have read and agreed to the published version of the manuscript.

---

**Supplementary Figure 1.**
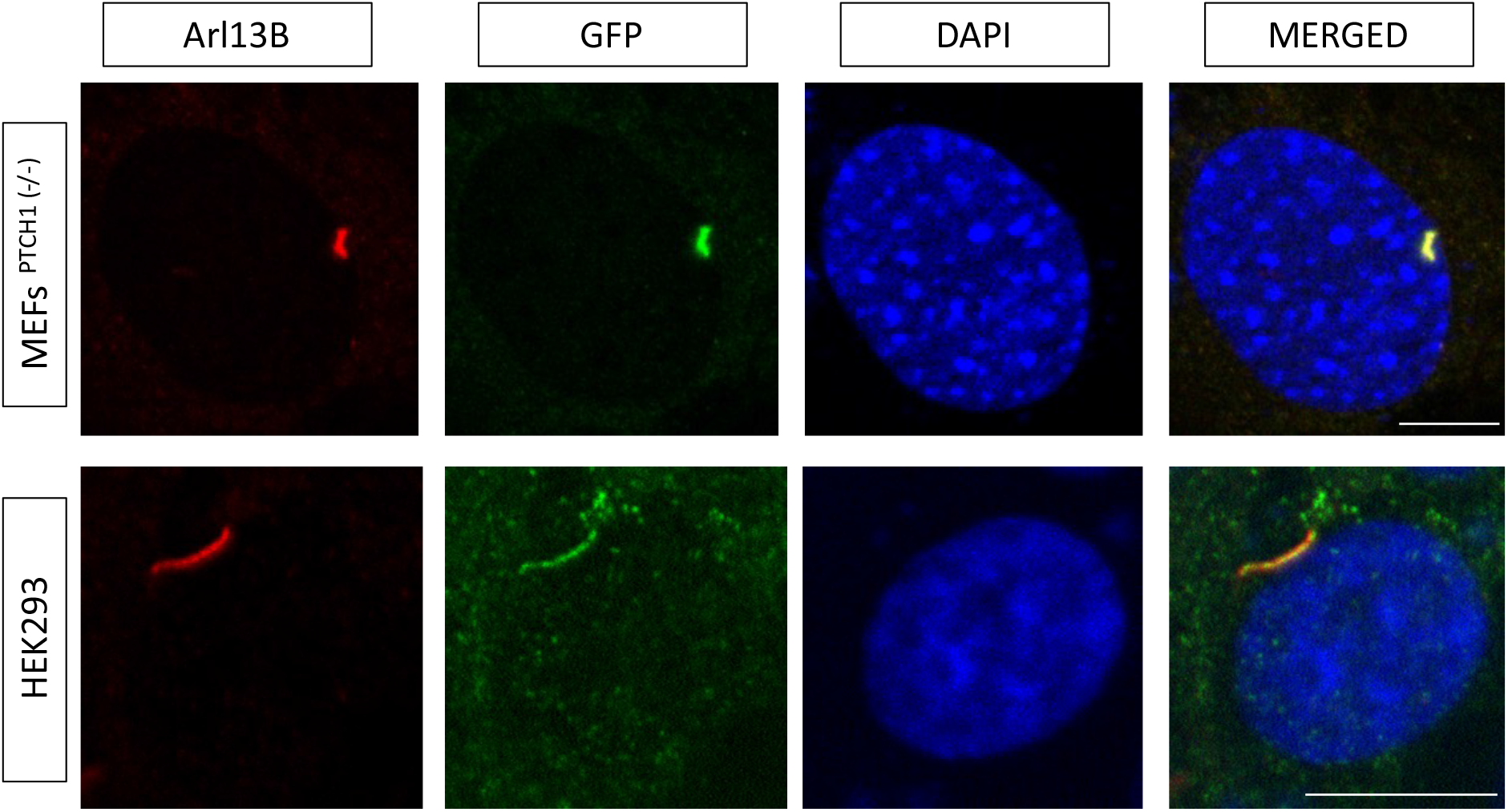
PTCH2-eGFP was transfected into *Ptch1*^−/−^ MEFs (top) or HEK 293 cells. After 24 h serum starvation to induce ciliogenesis, cells were fixed and stained with Arl13b (ciliary marker) and GFP, and then mounted with DAPI to counterstain nuclei. Scale bar = 10 μm.

## REFERENCES

1. Raleigh, D.R. & Reiter, J.F. Misactivation of Hedgehog signaling causes inherited and sporadic cancers. J. Clin. Invest. 129, 465–475 (2019).

2. Su, Y., Xing, H., Kang, J., Bai, L. & Zhang, L. Role the hedgehog signaling pathway in rheumatic diseases: An overview. Front. Immunol. 13, 940455 (2022).

3. Robbins, D.J., Fei, D.L. & Riobo, N.A. The Hedgehog signal transduction network. Sci. Signal. 5, re6 (2012).

4. Zhang, Y. et al. Structural Basis for Cholesterol Transport-like Activity of the Hedgehog Receptor Patched. Cell 175, 1352–1364 (2018).

5. Qi, C., Di Minin, G., Vercellino, I., Wutz, A, & Korkhov, V.M. Structural basis of sterol recognition by human hedgehog receptor PTCH1. Sci. Adv. 5(9), eaaw6490 (2019).

6. Kinnebrew, M. et al. Patched 1 reduces the accessibility of cholesterol in the outer leaflet of membranes. Elife 10, e70504 (2021).

7. Qian, H. et al. Inhibition of tetrameric Patched1 by Sonic Hedgehog through an asymmetric paradigm. Nat. Commun. 10, 2320 (2019).

8. Rudolf, A.F. et al. The morphogen Sonic hedgehog inhibits its receptor Patched by a pincer grasp mechanism. Nat. Chem. Biol. 15, 975–982 (2019).

9. Mann, R.K. & Beachy, P.A. Novel lipid modifications of secreted protein signals. Annu. Rev. Biochem. 73, 891–923 (2004).

10. Tukachinsky, H., Petrov, K., Watanabe, M. & Salic, A. Mechanism of inhibition of the tumor suppressor Patched by Sonic Hedgehog. Proc. Natl. Acad. Sci. U S A. 113, E5866–E5875 (2016).

11. Petrov, K., Wierbowski, B.M., Liu, J. & Salic, A. Distinct Cation Gradients Power Cholesterol Transport at Different Key Points in the Hedgehog Signaling Pathway. Dev. Cell 55, 314–327 (2020).

12. Kinnebrew, M. et al. Patched 1 regulates Smoothened by controlling sterol binding to its extracellular cysteine-rich domain. Sci. Adv. 8, eabm5563 (2022).

13. Myers, B.R., Neahring, L., Zhang, Y., Roberts, K.J. & Beachy, P.A. Rapid, direct activity assays for Smoothened reveal Hedgehog pathway regulation by membrane cholesterol and extracellular sodium. Proc. Natl. Acad. Sci. U S A. 114, E11141–E11150 (2017).

14. Goodrich, L.V., Milenković, L., Higgins, K.M. & Scott, M.P. Altered neural cell fates and medulloblastoma in mouse patched mutants. Science 277, 1109–1113 (1997).

15. Lee, Y. et al. Patched2 modulates tumorigenesis in patched1 heterozygous mice. Cancer Res. 66, 6964–6971 (2006).

16. Adolphe, C. et al. Patched 1 and patched 2 redundancy has a key role in regulating epidermal differentiation. J. Invest. Dermatol. 134, 1981–1990 (2014).

17. Zhulyn, O., Nieuwenhuis, E., Liu, Y.C., Angers, S. & Hui, C.C. Ptch2 shares overlapping functions with Ptch1 in Smo regulation and limb development. Dev. Biol. 397, 191–202 (2015).

18. Alfaro, A.C., Roberts, B., Kwong, L., Bijlsma, M.F. & Roelink, H. Ptch2 mediates the Shh response in Ptch1−/− cells. Development 141, 3331–3339 (2014)

19. Taipale, J. et al. Effects of oncogenic mutations in Smoothened and Patched can be reversed by cyclopamine. Nature 406, 1005–1009 (2000).

20. Chen, X.L. et al. Patched-1 proapoptotic activity is downregulated by modification of K1413 by the E3 ubiquitin-protein ligase Itchy homolog. Mol. Cell. Biol. 34, 3855–3866 (2014).

21. Li, X., Wang, J., Coutavas, E., Shi, H., Hao, Q. & Blobel, G. Structure of human Niemann-Pick C1 protein. Proc. Natl. Acad. Sci. U S A 113, 8212–8217 (2016)

22. Ohgami, N., Ko, D.C., Thomas, M., Scott, M.P., Chang, C.C. & Chang, T.Y. Binding between the Niemann-Pick C1 protein and a photoactivatable cholesterol analog requires a functional sterol-sensing domain. Proc. Natl. Acad. Sci. U S A 101, 12473–12478 (2004)

23. Pettersen, E.F. et al. UCSF Chimera - A visualization system for exploratory research and analysis. J. Comput. Chem. 25, 1605–1612 (2004).

24. Carpenter, D. et al. Characterization of two patched receptors for the vertebrate hedgehog protein family. Proc. Natl. Acad. Sci. U S A. 95, 13630–13634 (1998).

25. Pathi, S. et al. Comparative biological responses to human Sonic, Indian, and Desert hedgehog. Mech. Dev. 106, 107–117 (2001).

26. Nikaido, H. & Takatsuka, Y. Mechanisms of RND multidrug efflux pumps. Biochim. Biophys. Acta. 1794, 769–781 (2009).

27. Lu, X., Liu, S. & Kornberg, T.B. The C-terminal tail of the Hedgehog receptor Patched regulates both localization and turnover. Genes Dev. 20, 2539–2351 (2006).

28. Klenotic, P.A., Moseng, M.A., Morgan, C.E. & Yu, E.W. Structural and Functional Diversity of Resistance-Nodulation-Cell Division Transporters. Chem. Rev. 121, 5378–5416 (2021).

29. Zhang, Y. et al. Hedgehog pathway activation through nanobody-mediated conformational blockade of the Patched sterol conduit. Proc. Natl. Acad. Sci. U S A. 117, 28838–28846 (2020)

30. Rahnama, F., Toftgård, R. & Zaphiropoulos, P.G. Distinct roles of PTCH2 splice variants in Hedgehog signalling. Biochem. J. 378, 325–334 (2004).

31. Fan, Z. et al. A missense mutation in PTCH2 underlies dominantly inherited NBCCS in a Chinese family. J. Med. Genet. 45, 303–8 (2008).

32. Altaraihi, M., Wadt, K., Ek, J., Gerdes, A.M. & Ostergaard, E. A healthy individual with a homozygous *PTCH2* frameshift variant: Are variants of *PTCH2* associated with nevoid basal cell carcinoma syndrome? Hum. Genome Var. 6, 10 (2019).

